# Sensitivity analysis of genome-scale metabolic flux prediction

**DOI:** 10.1101/2022.07.30.502133

**Authors:** Puhua Niu, Maria J. Soto, Shuai Huang, Byung-Jun Yoon, Edward R. Dougherty, Francis J. Alexander, Ian Blaby, Xiaoning Qian

**Affiliations:** Texas A&M University, University of Washington at Seattle, Brookhaven National Laboratory, Lawrence Berkeley National Laboratory

**Keywords:** Sensitivity analysis, Bayesian network (BN) structure learning, regulated metabolic network modeling, metabolic engineering, optimal experimental design (OED)

## Abstract

TRIMER, Transcription Regulation Integrated with MEtabolic Regulation, is a genome-scale modeling pipeline targeting at metabolic engineering applications. Using TRIMER, regulated metabolic reactions can be effectively predicted by integrative modeling of metabolic reactions with Transcription Factor (TF)-gene regulatory network (TRN), where the TRN is modeled via Bayesian network (BN). In this paper, we focus on sensitivity analysis of metabolic flux prediction considering potential model uncertainty in TRIMER. We propose a computational strategy to construct the uncertainty class of TRN models based on the inferred regulatory order uncertainty when learning from given transcriptomic expression data and analyze the prediction sensitivity of the TRIMER pipeline for the metabolite yield of interest. The obtained sensitivity analyses can provide a useful guidance for Optimal Experimental Design (OED) to help acquire new data that can enhance TRN modeling and effectively achieve specific metabolic engineering objectives, including metabolite yield alterations. We have performed simulation experiments to demonstrate the effectiveness of our developed sensitivity analysis strategy and its potential to effectively guide OED.

**ACM Reference Format:** Puhua Niu, Maria J. Soto, Shuai Huang, Byung-Jun Yoon, Edward R. Dougherty,, Francis J. Alexander, Ian Blaby, Xiaoning Qian. 2018. Sensitivity analysis of genome-scale metabolic flux prediction. In *Proceedings of Make sure to enter the correct conference title from your rights confirmation email (CNB-MAC 2022)*. ACM, New York, NY, USA, 9 pages. https://doi.org/XXXXXXX.XXXXXXX

## 1 INTRODUCTION

Optimal Experimental Design (OED) and control with complex biological systems have significant impact on developing new computational strategies in systems and synthetic biology [3, 44] for targeted biochemical overproduction that may benefit human society, for example, in different energy-related and pharmaceutical applications [4, 5, 14, 16, 21, 22, 28]. In particular, metabolic engineering applies engineering principles to genetically engineer microbial strains by gene or reaction knockouts to optimize biological processes. Due to the demanding experimental cost and time to test different microbial strains *in vivo*, computational methods have been developed for *in silico* prediction of useful knockout strategies for beneficial mutants. However, many existing computational methods to obtain genetically engineered strains are based on genome-scale dynamic analysis at steady states, assuming the static network models [1, 2, 6, 13, 31–33, 38]. Recent efforts have integrated genetic regulatory relationships involving transcriptional factors (TFs) that may regulate metabolic reactions to achieve more accurate and robust prediction of target metabolic behavior under different conditions or contexts [8, 9, 11, 23, 25, 30, 34, 40]. To generalize these integrated hybrid models, we have developed Transcription Regulation Integrated with MEtabolic Regulation (TRIMER) [26] as a modeling pipeline targeting at metabolic engineering applications. Using TRIMER, regulated metabolic reactions can be effectively predicted by integrative modeling of metabolic reactions with TF-gene regulatory network (TRN), where the TRN is modeled via Bayesian network (BN) inferred from transcriptomic expression data. We have demonstrated promising metabolic flux prediction performances in both simulated and real-world microbial mutant design applications considering transcription regulation in genome-scale metabolic prediction [26, 27].

While all the existing efforts have demonstrated valid performances in selected model organisms with abundant data and careful manual curation, there is not much investigation on how model uncertainty, due to incomplete system knowledge and/or limited training data, may affect metabolic predictions. To the best of our knowledge, most of the existing works assume that the trained models are deterministic without considering potential model uncertainty. In this paper, we propose to analyze how model uncertainty may affect the metabolic engineering performance in our TRIMER framework, by considering an uncertainty class of learned network models instead of deterministic network models to investigate the metabolic engineering performance under uncertainty. In particular, through a mathematical programming formulation of Bayesian network topological ordering, we construct uncertainty classes of BN models for TRN and analyze the metabolic prediction sensitivity of our TRIMER modeling pipeline. We evaluate the sensitivity of the TRIMER pipeline by comparing the ground-truth metabolite yield alteration with TF knockout mutations with the predicted metabolic behavior based on the uncertainty class of perturbed BN models with different ranked network edge space based on BN topological ordering search. To be specific, we simulate both gene expression and metabolic flux data from a predefined ground-truth TRN-regulated metabolic network model, infer models from the simulated data and then construct uncertainty classes and check the prediction sensitivity by checking the correlation of the predicted and ground-truth metabolite yields. The obtained sensitivity analyses can provide useful guidance for model learning, calibration, and OED for metabolic engineering to allow biologists to better understand metabolism under perturbation and to take advantage of high-throughput genetic engineering for desired variants with reduced cost.

## 2 BACKGROUND

In [26, 27], we have developed an integrated regulatory-metabolic hybrid network model and genome-scale metabolic analysis pipeline, Transcription Regulation Integrated with MEtabolic Regulation (or TRIMER). TRIMER enables condition-dependent genome-scale metabolic behavior modeling and provides *in silico* predictions of metabolic engineering tasks, such as knockout phenotype and knockout flux predictions. As a hybrid network model, TRIMER has a metabolic network module that predicts metabolic fluxes based on the classic flux balance analysis (FBA) framework [10, 13, 20, 29, 38]. This well-known technique adopts Linear Programming (LP) with flux constraints introduced by metabolic regulation rules (i.e. a metabolic network) of a given organism, to model genome-scale steady-state metabolic status. For accurate and robust prediction of metabolic behavior under different conditions, regulatory relationships, in the corresponding transcription factor regulatory network (TRN) between transcriptional factors (TF) and their target genes, are integrated with the FBA formulation as additional transcriptional constraints over fluxes [1, 12, 18, 31, 33, 35]. Motivated by another genome-scale framework *PROM* (Probabilistic Regulation of Metabolism) [8], the authors of TRIMER have developed a TRN module that integrates the *a priori* known TF-gene interaction annotations and available gene/transcriptomic expression profiles to learn the corresponding Bayesian network (BN) for probabilistically modelling TRNs. The conditional probabilities *Pr*(*gene*(*s*)|*TF*(*s*)) can be inferred from the learned BN and used for the construction of regulatory constraints in the FBA formulation, predicting the metabolic behaviors under different conditions, for example TF knockouts in metabolic engineering. A typical formulation that predicts the regulated metabolic fluxes is shown below:

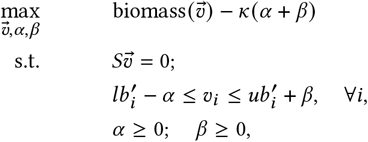

where *S* is the stoichiometric matrix deduced from the given metabolic network, 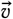 is a real-value vector representing metabolic fluxes of the hybrid network model in TRIMER, *α* and *β* can be considered as slack variables, *κ* is a hyper-parameter that controls the penalty of exceeding the fluxes bounds, *lb*^′^_*i*_ and *ub*^′^_*i*_ are regulatory flux upper/lower bounds computed based on *Pr*(*gene*(*s*)|*TF*(*s*)) and Flux Variability Analysis (FVA) [24]. Readers can refer to [26] for more details about flux bounds construction and how TRIMER predicts corresponding metabolic fluxes [27].

## 3 RESULTS

In this section, we first provide a brief overview of our proposed sensitivity analysis strategy. We then present the experimental results based on two simulated datasets to demonstrate the effectiveness of our proposed regulatory order based metabolic prediction sensitivity analysis.

### 3.1 Overview: Sensitivity analysis of TRIMER

Sensitivity analysis of TRIMER is achieved by evaluating metabolic flux prediction performances of the uncertainty class of BNs for TRN. In other words, we only focus on the model uncertainty of BN structures. To construct an uncertainty class, the essential idea is to infer the BN topological ordering from gene expression data and grow BN structures of various connectivity levels from the inferred node ordering. In this way, uncertainty classes of BNs at different perturbation levels can be constructed. By statistical analysis of metabolic predictions with different uncertainty classes, we can have better understandings on how TRN modeling may affect the final metabolic predictions in the TRIMER hybrid model. With such sensitivity analyses due to potential model uncertainty, practical guidance can be obtained to inform researchers for active learning and calibrating the modular components in TRIMER via optimal experiment design with the corresponding uncertainty quantification, for example defining the iterative structure updating policies for BN learning to improve metabolic prediction.

To develop such a sensitivity analysis capability in TRIMER, we first implement a BN topological order search algorithm to infer the network node ordering of BNs by formulating a mathematical programming optimization given transcriptomic expression data and the search space of BN edges. The topological ordering of BNs determines the parenthoods of nodes: an ancestor node must be of high orders than its descendant nodes. Therefore, given node parent sets and a network ordering of interest, the global optimal consistent structure can be obtained. To measure the uncertainty of the corresponding network ordering, ordering samples are obtained by bootstrapping [15] the adopted order search algorithms. Accordingly, the statistics of bootstrapped ordering samples can be computed to help derive the probability distributions of node ordering so that the uncertainty quantification can be derived.

The next question is how we may capture the potential model uncertainty from bootstrapped BN topological orderings and derive a rigorous mathematical framework to construct the classes of uncertain BN models for our proposed metabolic prediction sensitivity analysis. Following [39], we adopt a mathematical programming formulation to rank the pairwise orders of nodes corresponding to regulatory relationships in a given edge space. The formulation aims at identifying the critical regulatory relationships for which reducing the ordering uncertainty can help significantly improve the BN model fitting to the given training dataset. Solving this problem, all valid edges in the given edge space are ranked by their contribution to the uncertainty reduction, where the uncertainty is captured by the corresponding covariance matrix of ordering scores computed from bootstrapped samples. We then construct the BN uncertainty class of different size to complete the BN structure learning, where edges with higher rankings are more likely to be selected in the BN models.

Finally, transcription-regulated genome-scale metabolic prediction with the learned BNs in the corresponding uncertainty class can be done following TRIMER pipeline to investigate prediction sensitivity. The details of the TRIMER analysis pipeline can be referred to [27].

In summary, our proposed strategy is mainly comprised of three steps: 1) BN topological ordering, 2) Uncertainty class construction, and 3) metabolic prediction sensitivity analysis. Figure 1 provides a high-level overview of the strategy, depicting the main workflow.

**Figure 1:**
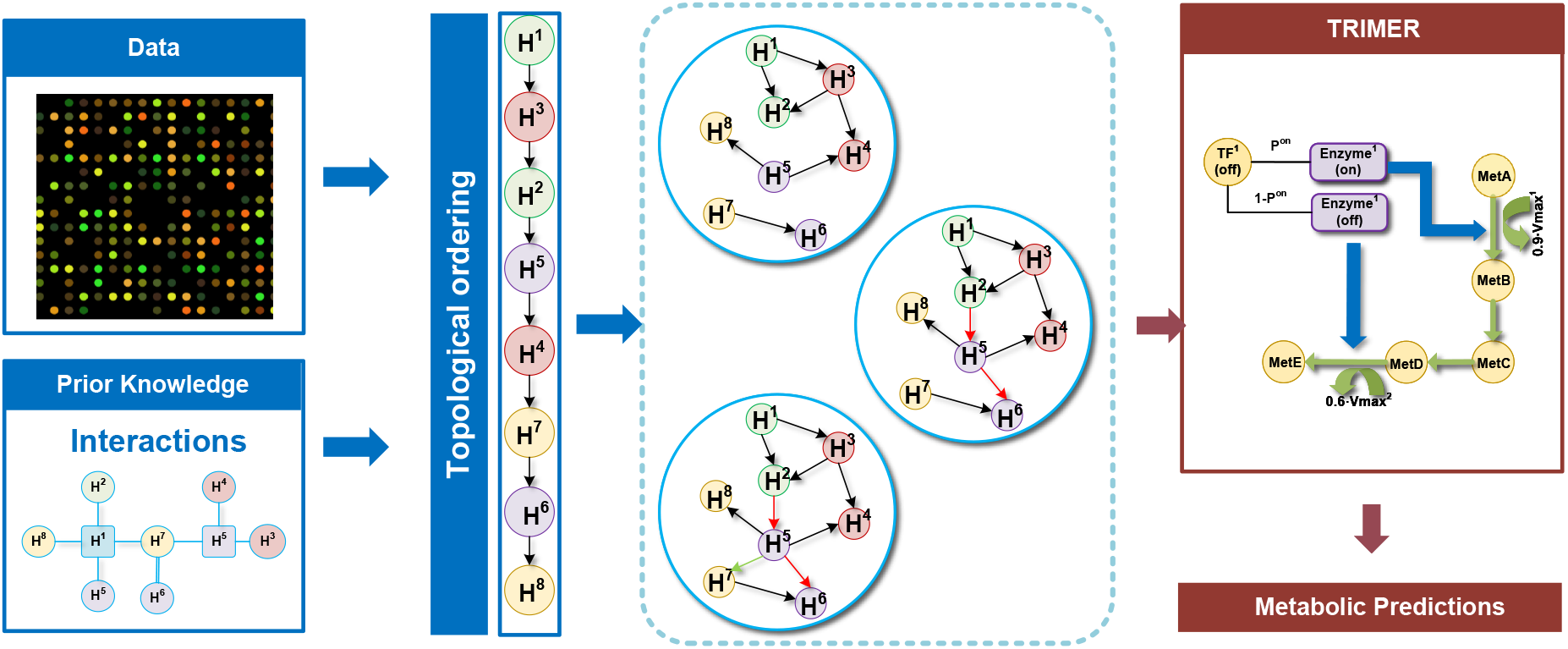
Illustrative workflow of the proposed sensitivity analysis strategy: 1) Gene expression data with the prior knowledge on regulatory interactions are used to infer the topological orderings of nodes in Bayesian networks (BNs). 2) The uncertainty class of BNs is constructed based on the uncertainty of the topological ordering due to incomplete knowledge and/or limited data. 3) TRIMER pipeline can be used with the uncertainty class for sensitivity analysis of the metabolic prediction.

### 3.2 Sensitivity analysis results

In our experiments, simulated ground-truth TRIMER models are used to generate gene expression data as well as metabolic flux data [26], which are used as the training datasets for BN learning and the ground-truth metabolic flux predictions for sensitivity analysis and performance evaluation. We focus on biomass prediction under multiple TF-knockouts while the proposed methods can be used for other metabolite yield predictions based on the problems of interest. We here use the two same simulated TRIMER models for *E. coli* with iAF1260 [19] as their genome-scale metabolic network model as described in [26]. One model is based on a small-scale TRN and the other is based on a large-scale TRN as detailed in the following subsections. More detailed descriptions on the TRIMER models, data sources, software requirements, hardware setups, as well as run-time estimation of each TRIMER component can be found in [26] and [27]. All the reported run-time in this paper is based on the implementation on a PC with Intel Intel i7 processor and 16GB RAM.

#### 3.2.1 Small-scale model sensitivity analysis

For the small-scale model, its corresponding TF-gene regulatory network (TRN) contains 50 nodes (12 TFs and 38 regulated genes) with 118 randomly generated edges. It is assumed that regulated genes can not be the parents of TFs, so the edge space contains 588 edges in total. Metabolic ground-truth in this model is simulated for all the 12 TF single-knockout conditions. Besides, we generate a gene expression dataset of 1000 samples from the ground-truth model. We bootstrap the Tabu order-based search for 100 times over the dataset obtaining 100 ordering samples, which can finish around 10 hours. The ordering deduced from the numerical mean of the bootstrapped orderings will be used as the base from which the ranked edge sets in the final BN uncertainty class are constructed. As the edge space is relatively small in this example, the edge ranking can be completed in a few seconds. The corresponding covariance matrix is shown in the left-hand side of Figure 2. Edges in the predefined edge space are ranked by their contribution to uncertainty reduction during the ordering posterior covariance updates via the mathematical programming formulation detailed in *Materials and Methods*. Next, uncertainty classes of Bayesian network structures consistent with the mean ordering are constructed from sampled edge sets with sizes ranging from 10% edge space to the whole edge space. Finally, the TRIMER model with TRN uncertainty classes is built and biomass predictions are made following the TRIMER pipeline. In our experiment, we calculate Pearson correlation coefficients (PCCs) between predictions and simulated ground-truths to evaluate TRIMER’s performance under a specific TRN configuration in the uncertainty class. The final metabolic prediction performance by TRIMER with different TRN uncertainty classes is illustrated in the left-hand side of Figure 3.

**Figure 2:**
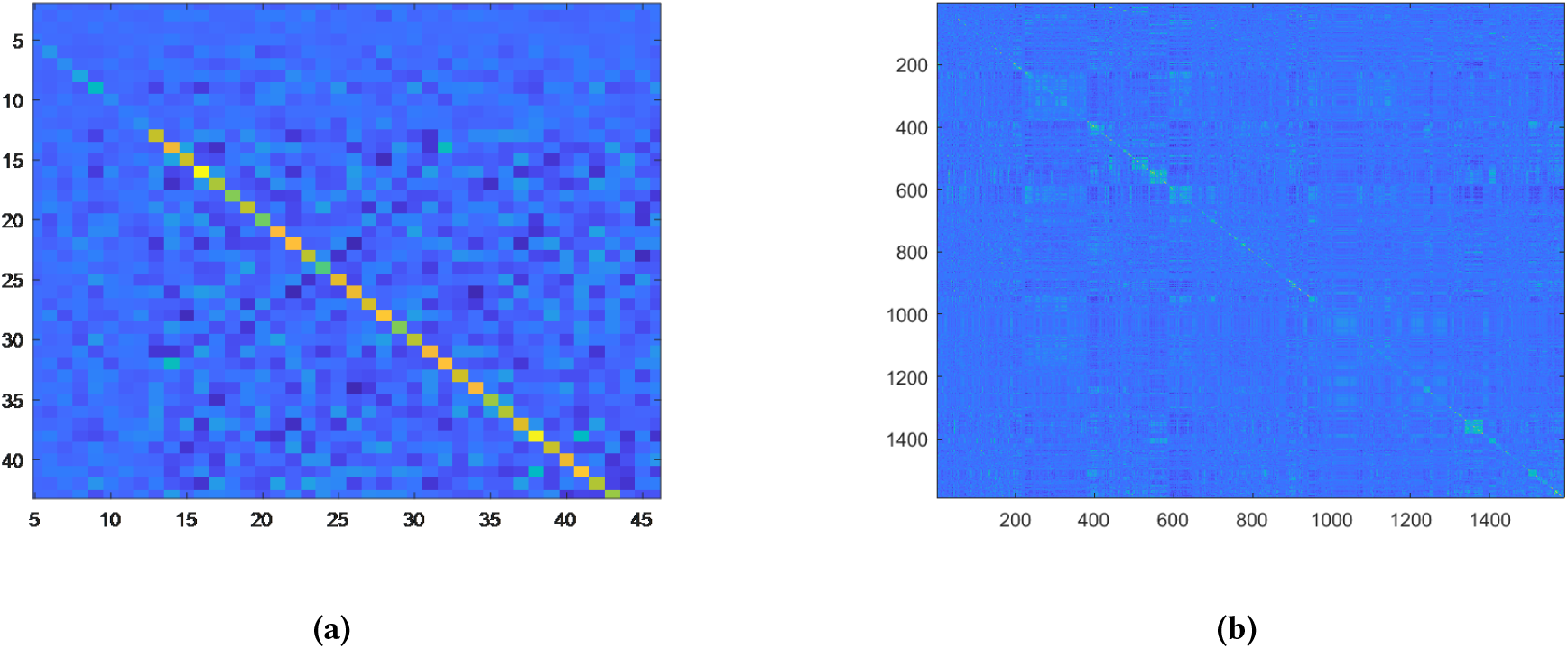
Estimated covariance matrices of bootstrapped ordering samples for (a) small-scale and (b) large-scale network models to help rank the pairwise TF-gene orders to construct the uncertainty class of BNs. Warmer colors indicate higher uncertainty regarding the pairwise ordering, hence specifying these edges may help significantly reduce the BN model uncertainty.

**Figure 3:**
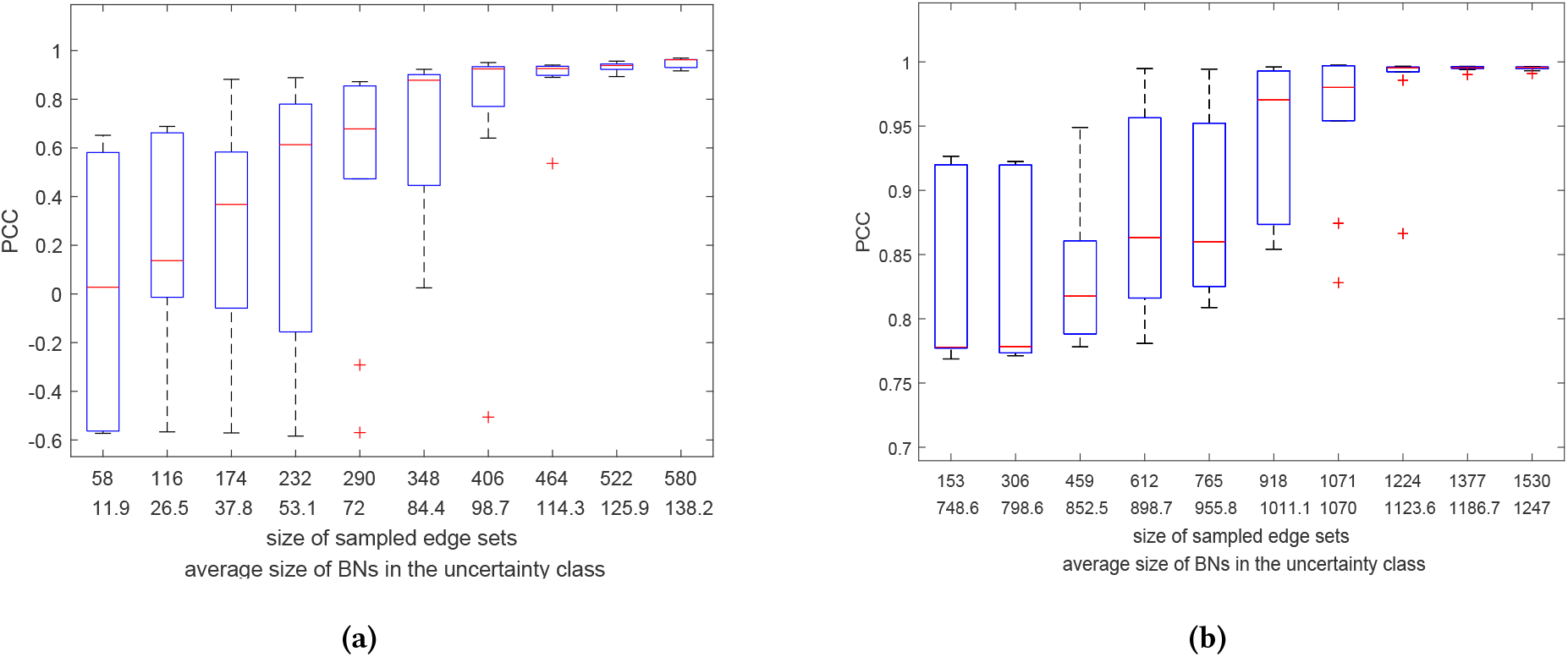
Box plots of Pearson correlation coefficient (PCC) between the metabolic prediction by TRIMER and the ground truth. Results are shown for (a) small-scale and (b) large-scale BN uncertainty classes as a function of the number of sampled edges and the average size of the BNs in the uncertainty class. The metabolic prediction performance improves when the uncertainty class is constructed from a larger edge space. Furthermore, the metabolic prediction sensitivity, illustrated by the quantile bars in the plot, decreases in general as the additional edges included in the edge space are less critical to achieve robust predictions.

From the plot, one obvious trend is when the uncertainty class is constructed with the allowed BN edge space covering more than 70% of the complete edge space, biomass predictions for TF-knockouts closely approach the simulated ground-truth results with the average PCC higher than 0.95. In general, the prediction performance improves when considering larger BN structure space as expected. More critically, the prediction sensitivity due to the model uncertainty decreases with the increasing edge search space. With the sampled ranked topological orderings, it is clear that the corresponding edges ranked in the lower 30% may not have much influence in both model prediction and sensitivity performances. On the other hand, when the constructed BN models miss highly ranked edges, metabolic predictions can be quite sensitive to potential model uncertainty even if their model predictions are satisfactory, as demonstrated when the uncertainty class construction covers the top ranked 40% to 70% edges of the complete edge space.

To further evaluate the prediction sensitivity of BN models for the small-scale TRIMER model, we conduct an experiment to investigate how its performance varies by changing the size of training gene expression datasets to construct the BN uncertainty classes. We have generated five expression datasets with sizes ranging from 200 to 1000. For each dataset, we evaluate the performance of TRIMER with the uncertainty classes constructed from the corresponding sampled edge sets whose sizes are fixed to the half of the complete edge space. The experimental results are depicted in Figure 4. Overall, with the increasing training samples for BN learning, both the average prediction accuracy, measured by PCC, and sensitivity, shown in the quantile bar plot, improve in general. The prediction performance may not improve much in average when we have more than 400 training samples. However, when investigating prediction sensitivity to the inherent model uncertainty with our regulatory order based uncertainty class construction, our results indicate that more training samples may need to guarantee the desired robust predictions. In our experiment, we observe that we require 1000 gene expression samples to achieve accurate and stable predictions.

**Figure 4:**
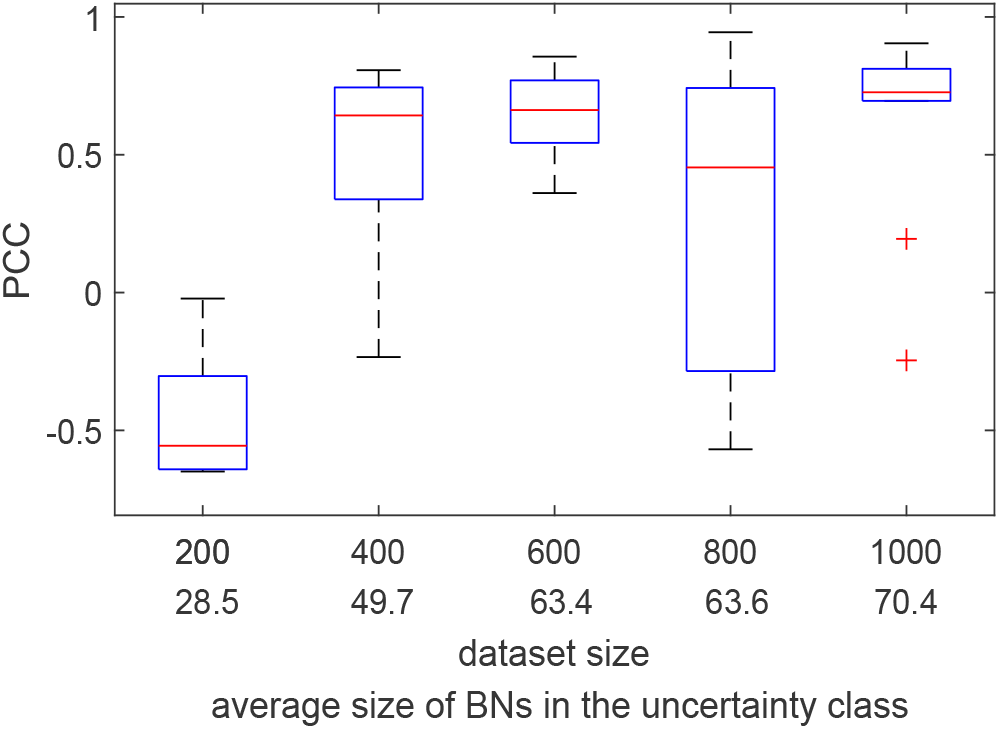
Box plots of Pearson correlation coefficient (PCC) between the metabolic prediction by TRIMER and the ground truth. Results are shown for uncertainty classes constructed based on simulated gene expression datasets of different sizes.

#### 3.2.2 Large-scale model sensitivity analysis

For the large-scale model, we consider a genome-scale TRN with 1509 edges randomly selected from the edge space containing 3704 edges in the annotated interaction list for *E. coli* in EcoMAC [7]. When constructing the BN uncertainty classes in this experiment, we only focus on an edge subspace comprised of 1533 edges, which connect genes directly regulating the reactions involving biomass production in the iAF1260 metabolic network model. We obtain 100 ordering samples by bootstrapping the order search over the simulated gene expression dataset of 1000 samples, similarly as described in the small-scale model experiment. In this set of experiments, the cyclic graph deduced from the interaction list is close to an acyclic graph with only 20 SCCs containing more than two nodes, for which Tabu order-based search can be completed within a minute. The corresponding covariance matrix is shown in the right-hand side of Figure 2. To construct the BN uncertainty classes, we first fix the BN structure for the genes that are not associated with biomass-related reactions to the optimal structure consistent with the derived mean topological ordering. The run-time of edge ranking for this larger graph is around ten minutes. We then sample different sizes of edge sets from the focused edge subspace. The corresponding BN edge space for the uncertainty classes also grows from containing 10% of the focused edge subspace to the complete space under consideration. The right-hand side of Figure 3 illustrates the performance under uncertainty.

We observe similar trends as in the small-scale model sensitivity analysis. In general, both metabolic prediction and sensitivity performances improve with the growing edge space. Due to the integration of the EcoMAC interaction list as prior knowledge for defining the BN edge space, we can achieve satisfactory prediction performance, PCC*>*0.95, when we cover more than 50% of the focused edges. On the other hand, to achieve stable predictions, we may need to cover more than 70% of the defined edge space.

## 4 MATERIALS AND METHODS

In this section, we provide a detailed description of our proposed sensitivity analysis strategy.

### 4.1 Order-based Tabu search

In the TRIMER pipeline described in [26], the Bayesian network structure learning is based on fitting the given gene expression profiles, denoted as *D*, to derive the conditional probabilities for a gene set *X* = {*X*_*n*_, *n* ∈ [1, *N*]} with a TF-gene interaction list denoted as *E* to capture the regulatory relationships of genes by their statistical dependencies, where *X* and *E* are interpreted as the set of network nodes and the search space of network edges. To learn the topological ordering of *X*, our implementation follows the same idea in [37], where an order-based heuristic search was proposed. Based on the score functions for candidate BN structures, the score of a given ordering ≺ can be defined as:

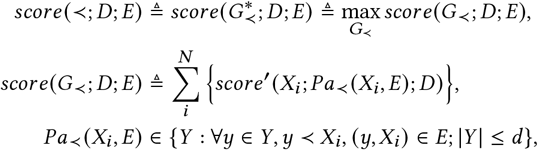

where *X*_*i*_ ∈ *X* denotes a node with its parent node set *Pa*_≺_ (*X*_*i*_, *E*) consistent with the ordering ≺ in the given edge space *E, y* ≺ *X*_*i*_ indicates that the order of *y* is higher than *X*_*i*_, *d* is the predefined upper bound of node’s in-degree, and *score*(*G*; *D*; *E*) is a decomposable score function used in the BN structure learning, such as Bayesian Information Criterion (BIC) score or BD (Bayesian Dirichlet) score. To find the best ordering, we use a heuristic search algorithm– multi-restart *Tabu* search—originally proposed in [17] based on the decomposed node-wise score function *score*^′^ (·;·;·). To perform the ordering search, we define a *swap* operator 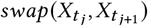 over nodes with the adjacent ordering in the *t*_*th*_ iteration as:

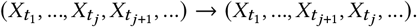

The best swapping operation is selected among all *n* − 1 candidate successors in the *t*_*th*_ iteration. This simplified search procedure based on the *swap* operator with significantly reduction of the ordering search space for computational efficiency of score comparison. Supposing that ≺^*t*^ is changed to a new ordering ≺^*t*+1^ by 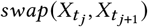, the delta-score of induced orderings only depends on the delta-score of 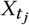 and 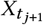. In addition, the only new operators deduced from ≺^*t*+1^ are just 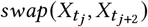 and 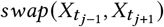. In each iteration, we can find the optimal parent set for any node in 𝒪 (*f*_*max*_)=𝒪(*N*^*d*^) as there are 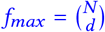 possible parent sets per node with the maximum in-degree *d* [37]. With *c* iterations, the time complexity of the algorithm is 𝒪(*cN*^*d*^). The pseudo-code of the implementation is provided in Algorithm 1.

#### Algorithm 1

Order-based Tabu search

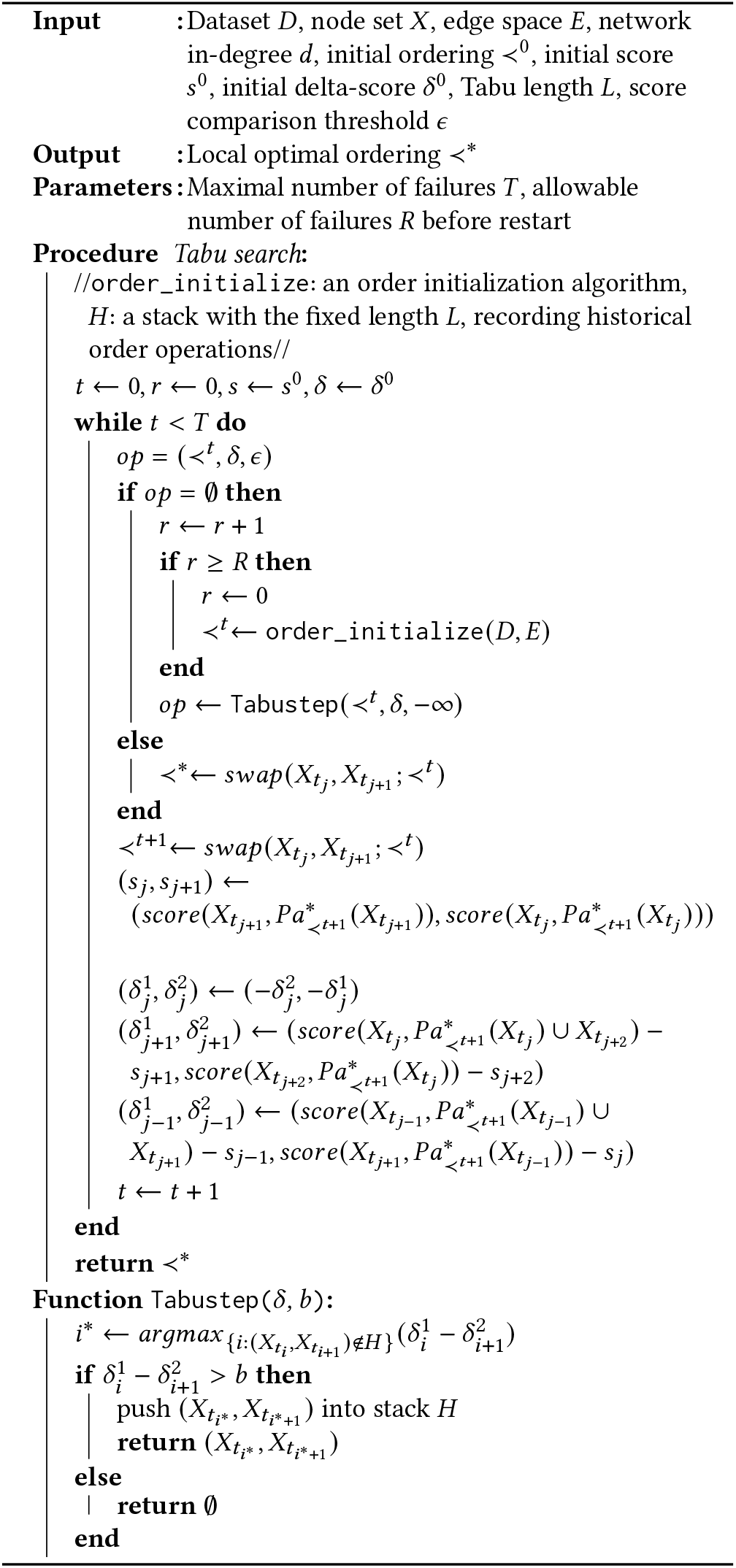

### 4.2 Order initialization

Initialization is crucial to guarantee satisfactory performance of heuristic search as the exact optimality is not ensured due to both the constrained search space and the greedy nature. In the extreme case, the given edge space corresponds to a directed acyclic graph (DAG), for which the optimal Bayesian network topological ordering is just any ordering consistent with the graph structure. However, in practice, the edge space almost always corresponds to a directed graph with cycles. While the node ordering can not be fully determined, it is still possible to obtain partial knowledge about it. For a directed graph *G* = (*X, E*), its strongly connected components (SCCs) denoted as *C* = {*C*_*m*_ : *m* ∈ [1, *M*]} are defined to be the maximal sets of nodes such that for each set, every pair of nodes within the set are reachable from each other. A graph of *C* can be denoted as *G*^*c*^ = (*C, E*^*c*^), where an edge exists between two SCCs if there is at least one edge between two nodes belonging to the two SCCs respectively. By the definition of the SCC, *G*_*c*_ must be a DAG. In cases when *G* is a DAG, *G*_*c*_ is just identical to *G*. Therefore, *G*_*c*_ determines a component-wise ordering ≺^*c*^, which provides us partial knowledge about how to initialize the node ordering ≺^0^ to inform the following order-based heuristic search algorithm. The globally optimal node-wise ordering ≺^*^ must be consistent with the ≺^*c*^ ordering while relative orders among nodes within the same SCC is still undetermined. To obtain an appropriate initial node ordering ≺^0^, *G*_*c*_ and the corresponding ordering ≺^*c*^ are first identified from *G* = (*X, E*), where SCCs are identified by Tarjan’s algorithm [36] with the computational complexity 𝒪(|*E*| + |*X*|). Next, node ordering ≺^0^ consistent with ≺^*c*^ can be found easily, where relative orders of nodes belonging to the same SCC are randomly generated. The pseudo-code of the initialization algorithm is shown in Algorithm 2 and the corresponding workflow is illustrated in Figure 5.

**Figure 5:**
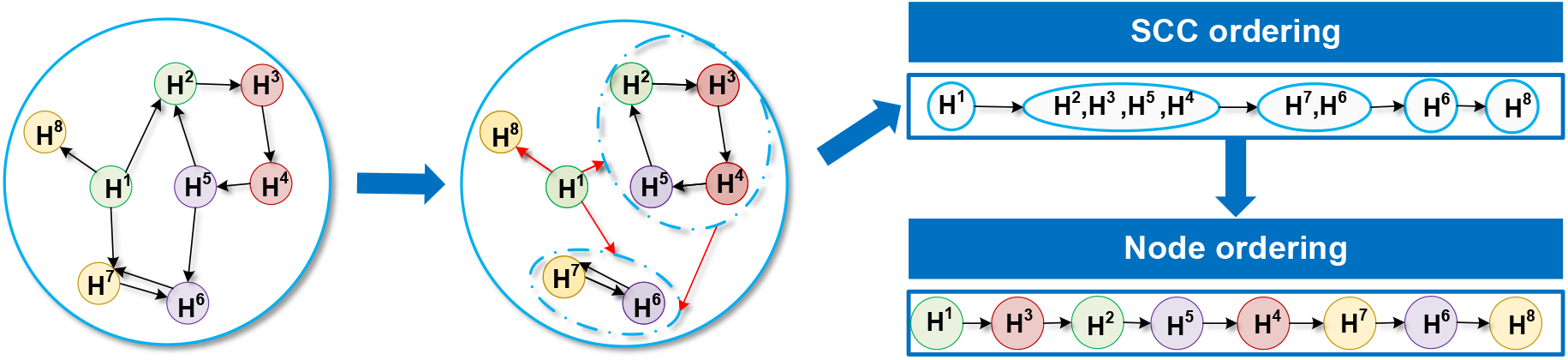
Illustration of the order initialization workflow. First, the directed acyclic graph (DAG) of strongly connected components (SCCs) is identified from the given directed graph; then a node-wise ordering consistent with the SCC ordering is randomly selected.

#### Algorithm 2

Order initialization from a given edge space

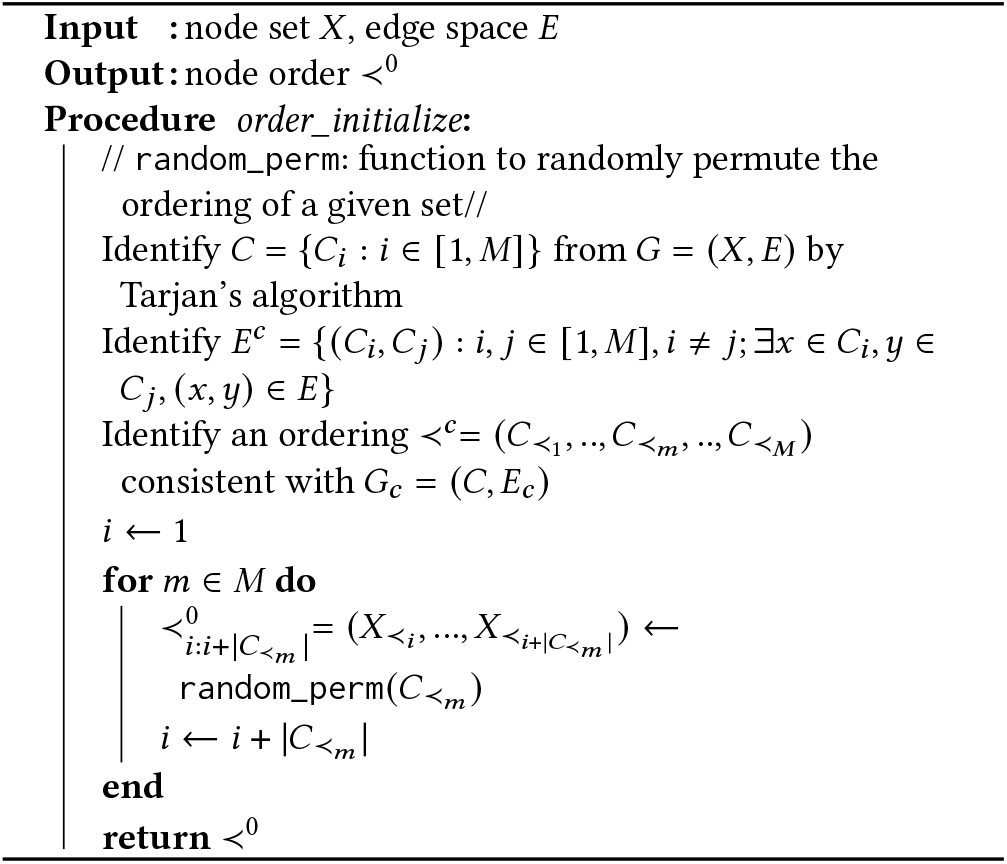

### 4.3 Semidefinite programming formulation for uncertainty class construction

First of all, we use a multivariate Gaussian random vector *ϕ* ∈ *R*^|*X*|^ ∼ 𝒩(*μ*, Λ^−1^) as the numerical representation of an ordering ≺, where the nodes of higher order are supposed to have larger values in *ϕ*. By *bootstrapping* the previously described order-based search algorithm, we can estimate the corresponding distribution parameters of 𝒩(*μ*, Λ^−1^) by the bootstrap samples of *ϕ. Bootstrapping* here means repeatedly perturbing *D* and applying the order search algorithm on the perturbed data sets to obtain a set of perturbed local optimal orderings. To establish the relationship between node-wise orders and edges, we associate edge *E*_*k*_ = (*X*_*i*_, *X*_*j*_), *E*_*k*_ ∈ *E* with a real-value random variable *y*_*k*_ ∼ 𝒩(*ϕ*_*i*_ − *ϕ*_*j*_, *γ*^−1^) to represent the pairwise order difference, where *γ* is a hyper-parameter. Values of *y*_*k*_ can indeed be interpreted as the confidence of the corresponding pairwise ordering: the larger the value of *y*_*k*_ is, the more confident we are to support the ordering induced by edge *E*_*k*_. As proposed in [39], a binary matrix *B* ∈ {0, 1}^|*E*|×|*X*|^ can be used to collectively represent all edges in *E*, in which *B*_*k,l*_ for *E*_*k*_ = (*x*_*i*_, *x*_*j*_) is defined as:

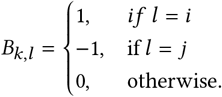

In cases when only a subset of *E* is considered, a binary matrix *B*^*^ = *Diag*(*v*)*B* is used as the corresponding matrix representation, where *v* ∈ {0, 1}^|*E*|^ is a binary vector, *Diag*(*v*) denotes the corresponding diagonal matrix. Therefore, *y* ∼ 𝒩(*B*^*^*ϕ*, Γ^−1^) represents the pairwise ordering confidence about all the edges of interest, where Γ is a hyper-parameter diagonal matrix. It should be pointed that *P*(*ϕ*|*μ*, Λ^−1^) is a conjugate prior of *P*(*y*|*B*^*^, *ϕ*, Γ^−1^) as they are both Gaussian. Therefore, it can be easily verified [39] that:

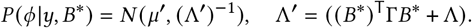

It can be observed that Λ = (*B*^*^)^T^Γ*B*^*^ quantifies the effect of the edge set of interest over the uncertainty of ordering *ϕ*. Therefore, a straightforward idea to rank edges is by their contribution to the uncertainty reduction. To be more specific, the ranking is achieved by solving a semidefinite programming problem (SDP) proposed in [39], which is defined as:

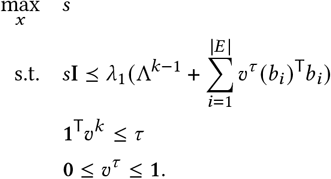

In the formulation above, the column vector *b*_*i*_ ∈ {0, 1}^|*X*|^ corresponds to the *i*_*th*_ row of matrix *B*, 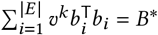 and *τ* is a scalar representing the size of a selected edge subset. Values of resulting *v*^*τ*^ represents the top *τ* edges in items of uncertainty reduction. By solving the SDP repeatedly for |*E*| times with *τ* increasing from 1 to |*E*|, the growing ordering of selected edges implies a ranking of edges in *E*. It should be pointed out that *v*^*τ*^ is relaxed to [0, 1]^|*X*|^ to guarantee the convexity of the SDP. While this relaxation leads to potential interpretation ambiguity of the solution as the number of non-zero elements in *v*^*τ*^ can be higher than *τ*. However, we can still select the top *τ* edges by the magnitudes of values in *v*^*τ*^. To construct uncertainty classes of BN models for TRN in TRIMER, ranked edges are assumed to comply with a distribution defined as follows:

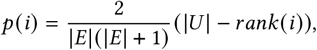

where *i* denotes the index of an edge in *E* and *rank*(*i*) denotes its rank in terms of uncertainty reduction. Then edge sample sets with various sizes can be drawn from this distribution. The corresponding structure *G*^*S*^ for an edge sample set *S* is identified by 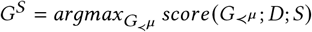, where ≺^*μ*^ is deduced from the numerical mean *μ* of the ordering samples.

### 4.4 TRIMER as a simulator

As describe in [26], TRIMER can serve as a simulator of gene expression and metabolic flux data. Given a ground-truth BN model for TRN with appropriate conditional probability tables for each node in TRN, gene expression datasets can be simulated by drawing sampling from the distribution described by the BN. Moreover, conditional probabilities with respect to TFs and target genes can be inferred from the BN and used for constructing regulatory flux constraints for genome-scale metabolic predictions when integrating with the available metabolic reaction network model. Adding these new constraints into the corresponding FBA formulation for the metabolic network of a given organism, condition-dependent metabolic states of the organism can be simulated and treated as the ground-truth metabolic fluxes.

For sensitivity analysis, we first simulate the ground-truth metabolic fluxes to estimate the biomass for different TF-knockout *E. coli* strains as described in [26]. We investigate the model uncertainty by computing the corresponding Pearson correlation coefficients (PCCs) between ground-truth biomass fluxes and the TRIMER predicted fluxes with uncertainty classes constructed based on the simulated gene expression data, of which the BN structures may significantly deviate from the ground-truth model that simulates the data.

## 5 CONCLUDING REMARKS

From our experimental results, it can be observed that when the BN models deviate more from the optimal BN by missing highly ranked critical edges, the prediction accuracy, measured by correlation between ground-truth biomass fluxes and predicted fluxes of the perturbed BN models, does decrease. More critically, when constructing such uncertainty model classes and investigating prediction sensitivity, our results indicate that we may need better prior knowledge and more training data to achieve both accurate and stable predictions. Our sensitivity analyses also indicates that reliable uncertainty quantification may require more data. While many existing works have reported valid performances in selected experiments, there may still be potential overfitting risks with predictions not easy to generalize when having slightly perturbed systems.

Our regulatory order based sensitivity analysis also helps identify the set of edges, for which the corresponding uncertainty reduction can significantly help model prediction and improve sensitivity. Such a capability can lead to new uncertainty quantification formulations, which may enable optimal experimental design strategies for active model learning [41–43] and more robust intervention strategies in metabolic engineering, which we leave for future research.

